# Impacts of a millennium drought on butterfly faunal dynamics

**DOI:** 10.1101/242560

**Authors:** Matthew L. Forister, James A. Fordyce, Chris C. Nice, James H. Thorne, David P. Waetjen, Arthur M. Shapiro

## Abstract

Climate change is challenging plants and animals not only with increasing temperatures, but also with shortened intervals between extreme weather events. Relatively little is known about diverse assemblages of organisms responding to extreme weather, and even less is known about landscape and life history properties that might mitigate effects of extreme weather. We find that northern California butterflies were impacted by a millennium-scale drought differentially at low and high elevations. At low elevations, phenological shifts facilitated persistence and even recovery during drought, while at higher elevations a shortened flight season was associated with decreases in species richness. Phenological and faunal dynamics are predicted by temperature and precipitation, thus advancing the possibility of understanding and forecasting biological responses to extreme weather.

## Main text

Extreme weather events have occurred with increasing severity and frequency in recent decades, a trend that has been linked to anthropogenic climate change and shifts in atmospheric circulation^1^. Human societies are vulnerable to disruption by weather phenomena that include hurricanes and heat waves, and these events add to and interact with other climate change effects^2^. Natural systems of plants and animals have been studied intensively from the perspective of shifting average climatic conditions^3,4^, but we know less about the impacts of either elevated variation or extreme events on wild organisms^5,6^. Here we take advantage of decades of data on 163 butterfly species across an elevational gradient in Northern California (Fig. 1a) to address knowledge gaps within the context of a severe, millennium-scale drought that impacted the region from 2011 to 2015^7^. Specifically, we asked the following: 1) does an extraordinary, multi-year drought elicit a faunal response that is extreme relative to faunal behavior in previous dry years? 2) Are impacts on the fauna consistent or divergent across elevations? A theoretical expectation is that organisms living in more heterogeneous environments should be more resilient to extremes of temporal variation^8^. We predicted that butterflies at montane sites would be robust relative to populations at lower elevations in landscapes that are both less spatially variable and already impacted by a history of human activity. 3) Finally, we asked if population-level responses to drought are mediated by phenological shifts. Species that are able to begin activity earlier in the spring might reach higher population densities^9^, potentially offsetting detrimental drought effects. Another possibility is that ectotherms exposed to longer growing seasons could fall into a developmental trap by which extra generations fail because of insufficient time^10^.

**Figure 1.**
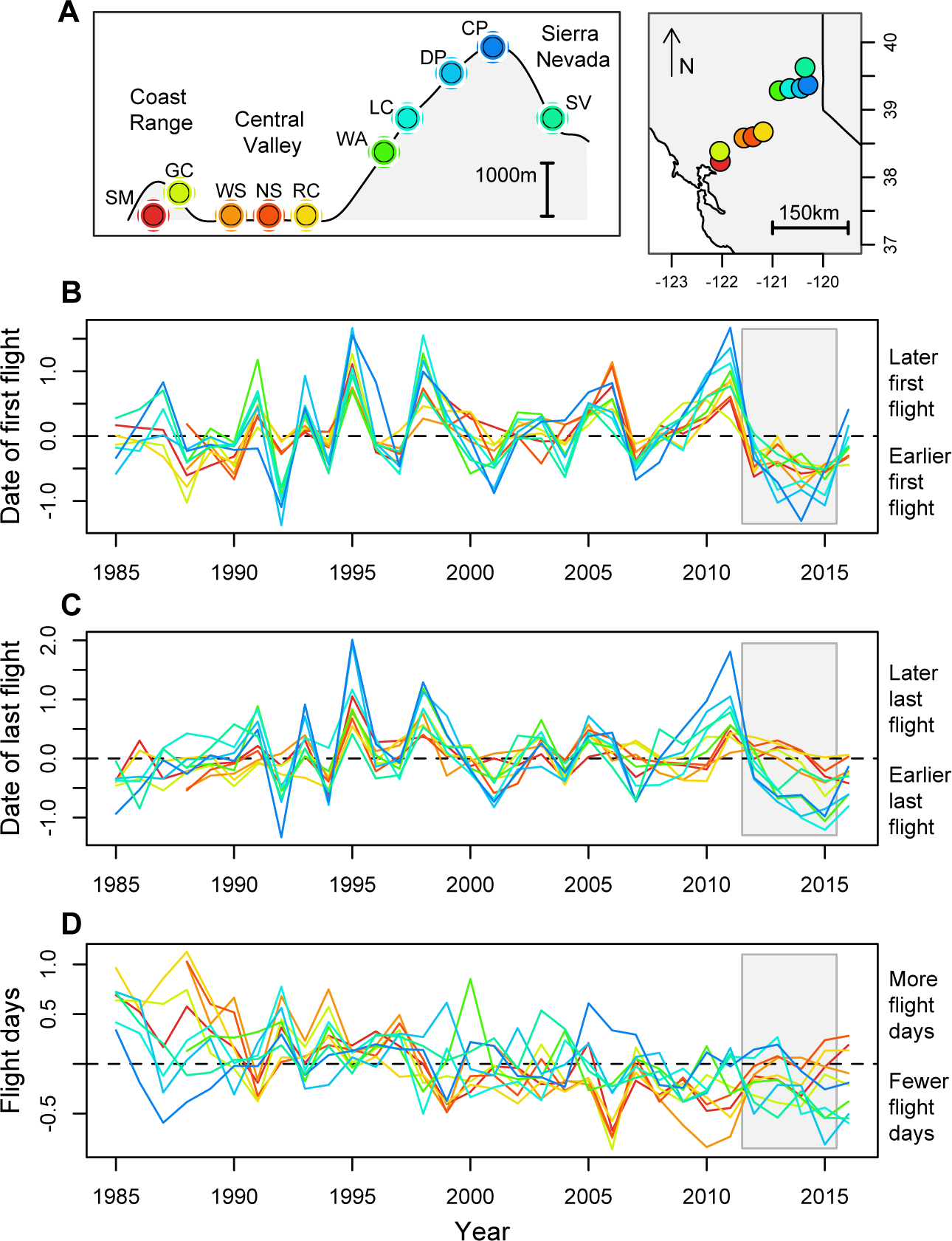
(**a**) Elevational profile of Northern California (left) and map of the same area (on the right) with ten study sites indicated on both; site names as follows, from west to east: Suisun Marsh (SM), Gates Canyon (GC), West Sacramento (WS), North Sacramento (NS), Rancho Cordova (RC), Washington (WA), Lang Crossing (LC), Donner Pass (DP), Castle Peak (CP) and Sierra Valley (SV). (**b**) Average dates of first flight and (**c**) last flight across species at each location and year. (**d**) Average flight days, which are the average fraction of days individual species are observed per year. In panels **b, c,** and **d**, color coding for individual lines corresponds to sites as in panel **a**, and the data are shown as *z*-standardized values. Grey rectangles in panels **b, c**, and **d**, indicate the major drought years from late 2011 into 2015.

Investigations of butterflies at our focal sites have reported that a majority of populations at the lowest elevations have been in decline since at least the mid-1990s^11^, which has been attributed to changes in land use and warming temperatures^12^. Populations at higher elevations, in contrast, have shown relatively less directional change over time, with the exception of a decline in more dispersive, disturbance-associated species that rely on demographic connections with lower-elevation source populations^13^. Previous analyses of abiotic effects, prior to the 2011–2015 drought, have noted responses to weather that were heterogeneous and idiosyncratic among sites and species^14,15^. In contrast to the previously-documented heterogeneity in population response, we find here that the recent, extreme drought years resulted in a number of faunal responses that were consistent across sites and elevational subsets of sites (montane versus valley sites).

## Results

A prolonged and consistent shift towards earlier spring flights can be seen in Fig. 1b. While the shift in phenology is evident across elevations, the dynamics of the flight window diverge later in the season: at higher elevations, the date of last flight shifted to an earlier time during the drought, while at lower elevations the last flight dates from 2011 to 2015 are closer to the long-term average (Fig. 1c). In other words, the total flight window expanded at lower elevations, while in the mountains the flight window shifted and compressed towards the start of the season, a change that is reflected in fewer overall flight days at higher sites (Fig. 1d).

Along with the recent reduction in the average number of days that butterflies were observed flying at higher elevations during the drought years, there have been fewer butterfly species observed per year at the same sites (Figs. 2a – 2e). In some cases, the millennium drought was associated with a discrete downturn (e.g., Figs. 2b and 2c), while at other montane sites the recent drought years contributed to ongoing, negative trends (Figs. 2d and 2e). A downward trend in species richness is less evident at the highest site (CP, Fig. 2a), which previous analyses have found to be receiving immigrants from lower elevations as populations shift upslope in warmer years^13^. In a dramatic reversal of fortunes, the lowest elevation sites during the millennium drought experienced some of their most productive years in nearly two decades, as reflected both in numbers of species (Figs. 2g – 2j) and numbers of individuals observed (Figs. 2l – 2o). Results shown in Fig. 2 are for simple richness (the number of species observed per year). We repeated the analyses using alpha diversity Hill numbers that down weight the importance of rare species (Fig. S1, S2), and found similar results for all sites except for GC, where a long-term decline in the number of species becomes evident when rare, or transient species have less influence.

**Figure 2.**
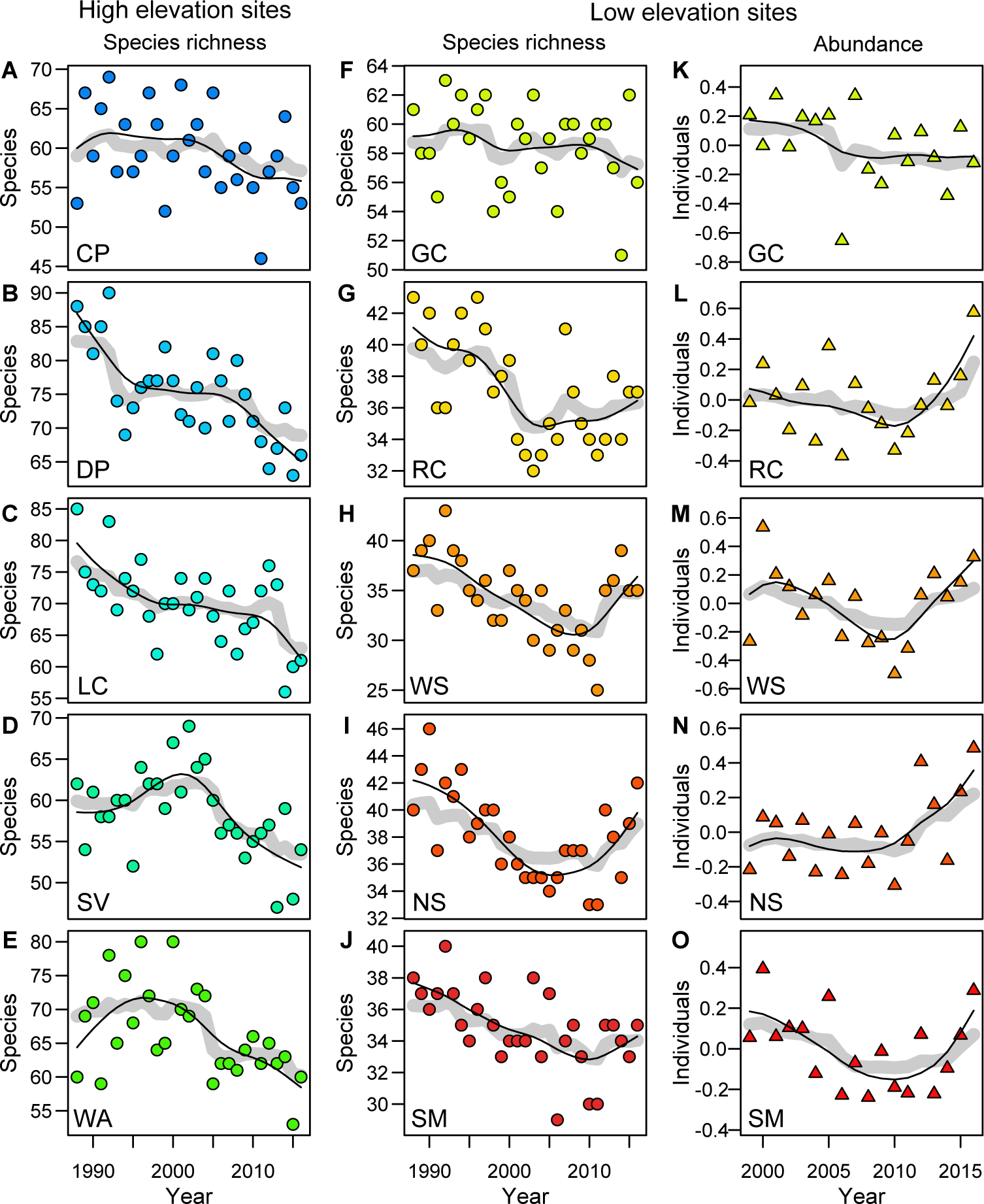
Species richness (**a** through **j**) for all sites, and abundance (**k** through **o**) for low elevation sites (abundance data was only collected at the low sites). Two letter site names and colors follow Fig. 1A. In all panels, patterns are visualized with both a spline fit with five degrees of freedom (thin black line) and predicted values from random forest analysis (thick gray line) incorporating variation in sampling effort. In panels **k** through **o**, values plotted are *z*-standardized values of total abundance (number of individuals) per year averaged across species.

Why did the low elevation sites apparently rebound during the drought years? Using the lowest sites (RC, WS, NS and SM) and a span of years starting just before the millennium drought, we discovered an effect of phenological plasticity. Specifically, species whose first flight shifted to an earlier day were the species that became more abundant (*r* = −0.50, *P* < 0.001; Fig. S3). Butterflies at the lowest elevations are almost entirely multivoltine, and an earlier start for those species led to an extension of the flight season (Fig. 1d), and an increase in abundance and richness (Fig. 2). To understand climatic drivers of phenology, at low and high elevations, we modeled the dates of first flight as a function of maximum and minimum temperatures, precipitation, and El Niño (ENSO) conditions. Models explained 60% of the inter-annual variation at low elevations *(F_6138_* = 35.17, *P* < 0.001), and 72% at the higher elevation sites *(F_6_,_138_* = 59.68, *P* < 0.001) (Table S1). Minimum and maximum temperatures had negative effects on first flight dates (warmer temperatures lead to earlier flights), and the effect of the former was most noticeable at higher elevations (Figs. 3a and 3b). Precipitation, as reflected by local weather and ENSO conditions, had a delaying effect on phenology (positive β coefficients in Figs. 3c and 3d), which is expected as wetter conditions are associated with cooler, cloudy days and delayed spring emergence.

**Figure 3.**
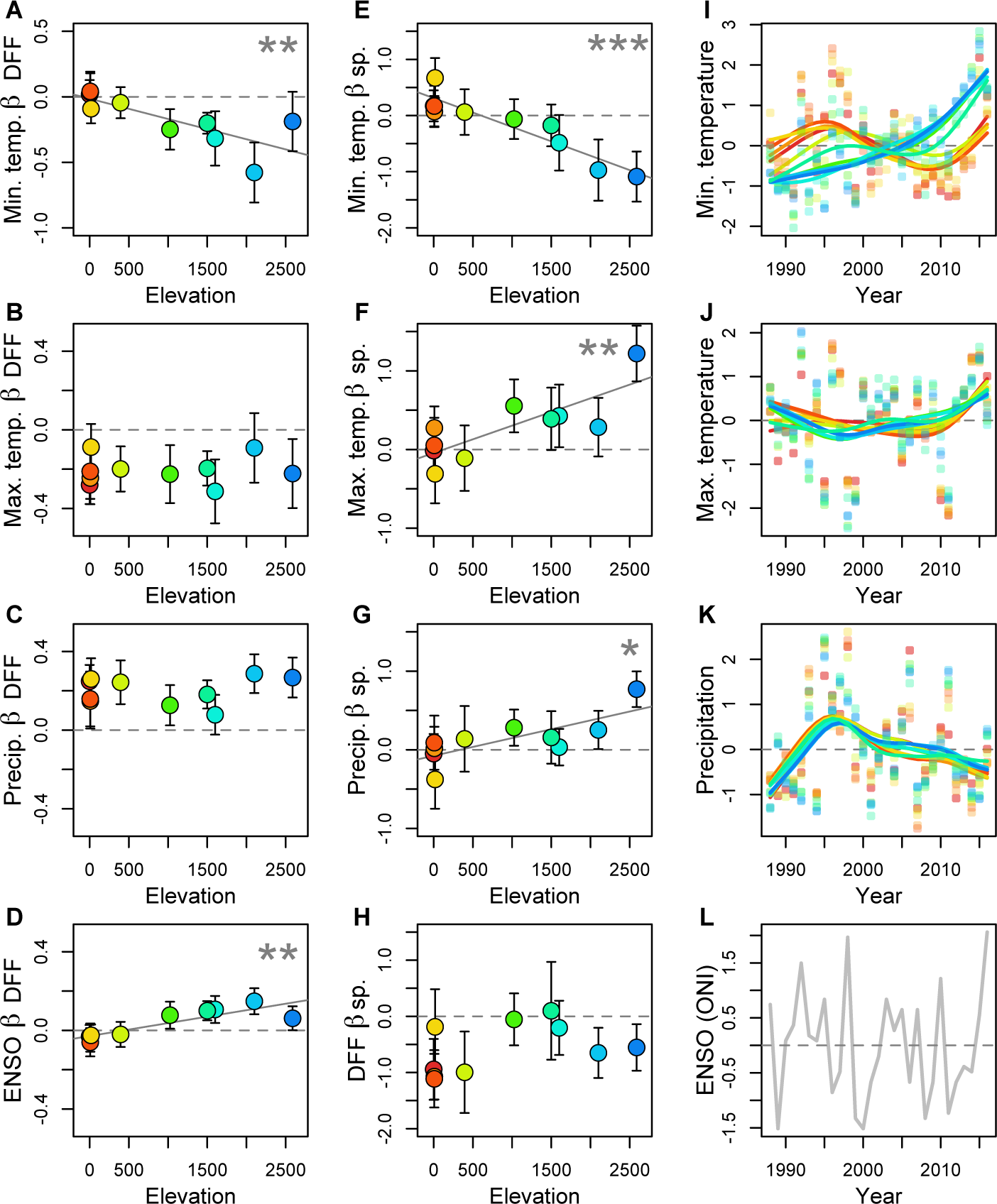
Results from a model of phenology (**a** through **d**) and a model of species richness (**e** through **h**), as well as plots of weather variables through time (**i** through **l**). In the model results (**a** through **h**), the values shown are β coefficients (with standard errors) that summarize the effect of a particular weather variable (while controlling for other variables) on either phenology or richness for each site. Trend lines are only shown in plots where the effect of a particular weather variable changes with elevation (* *P* < 0.05; ** *P* < 0.01; *** *P* < 0.001; see Table S4 for additional details). In panels **i** through **k**, weather patterns are visualized using splines with five degrees of freedom; panel **l** is the El Niño index (ONI) for each year. Weather variables are shown as *z*-standardized values in panels **i** through **k**, and site specific colors in all plots are the same as in Fig. 1a.

Models of species richness revealed even more pronounced variation in weather effects across elevations, including an increased importance of variation in minimum temperatures (Fig. 3e), maximum temperatures (Fig. 3f), and precipitation at higher elevations (Fig. 3g) (see Table S2 for full details). The highest elevations are most negatively affected by dry years with warmer nighttime temperatures. While daily maximum temperatures have risen everywhere (Fig. 3j) and patterns of precipitation have fluctuated in concert across sites (Fig. 3k), minimum temperatures have risen most steeply at the mountain sites (Fig. 3i). Models of species richness also included phenology (date of first flight) as an explanatory variable, and we found an overall negative association (Fig. 3h), such that earlier emergence is associated with elevated richness (consistent with the effect of phenology on abundance at low elevations; Fig. S3). However, the beneficial effects of earlier emergence at higher elevations might not be as consequential because of a lack of multivoltine species, or they may simply be outweighed by the negative effects of minimum temperatures. Negative effects of minimum temperatures at the higher elevation sites range from 0.48 fewer species seen for every degree Celsius of warming at WA, to 6.46 fewer species seen for every degree at DP (Table S3).

## Discussion

In contrast to previous observations that extreme weather elicits species-specific responses^16^, the millennium California drought produced consistent responses across many sites that included advanced dates of first flight with elevation-specific changes in flight windows and species richness. The resilience exhibited by the lowest elevations is associated with phenological flexibility combined with multivoltine life histories and climatic associations that are less detrimental (in the context of current climatic trends) than biotic-abiotic associations at higher elevations. Many researchers have hypothesized an impending mismatch between trophic levels as a result of climate change^17^. The results from low elevation butterflies in California perhaps challenge that hypothesis, or at least suggest that a shift in phenology at the consumer trophic level need not always have negative consequences. In addition to having multiple generations each year, populations at the lowest elevations have access to agricultural lands. Although association with irrigation does not appear to predict population dynamics during the drought (Fig. S3), we cannot rule out the possibility that low elevation populations were buffered during the drought by irrigated crops or agricultural margins. If true, a reliance on agriculture would be interesting in the light of a recent hypothesis that long-term declines in low elevations butterfly populations are associated with intensification of pesticide applications^18^. It is possible that the rebound of the drought years could be followed by a more severe decline following concentrated agriculture dependency and toxin exposure.

It has been known for some time that high latitude environments are warming faster than the rest of the planet. It is only recently that climatologists have become aware that higher elevations may also be experiencing a disproportionate share of warming^19^, which raises the issue of how cold-adapted, montane ecosystems will respond. Contrary to the expectation that mountains offer microclimatic refugia and preadapt species for climatic variation^8,20^, we found high elevation butterfly communities to be declining and especially sensitive to dry years with warmer minimum temperatures. Warmer and drier years are associated with lower productivity of mesic-adapted plant communities^21^, and shorter windows during which montane plants are optimal for nectar and herbivory^22^. We did not model snowfall because it is highly correlated with annual precipitation at our sites (see Methods), but reduced snowpack in dry, warm years would have additional negative effects including higher overwinter mortality for life history stages that typically spend the winter under a blanket of snow^23^. The daily temperature range (the difference between maximum and minimum temperatures) has been shrinking around the globe^24^, but the ecological consequences of this thermal homogenization are poorly understood and not yet incorporated into theoretical expectations of global change biology^25^. The results reported here suggest that we have much yet to learn about organismal responses to extreme weather at low and high elevations, but that powerful and simple models predicting faunal dynamics are possible for ectotherms even in the face of unprecedented climatic variation.

## Methods

### Butterfly data

Ten study sites (Fig. 1a) were visited biweekly (every two weeks) by a one of us (AMS) for between 45 and 29 years, depending on the site, and only during good “butterfly weather” when conditions were suitable for insect flight (nearly year round at low elevations, and during a more narrow period at higher elevations). At each site, a fixed route was walked and the presence and absence of all butterfly species noted. Maps of survey routes and site-specific details, as well as publically-archived data can be found at http://butterfly.ucdavis.edu/.

For most analyses, we restricted data to a common set of years, from 1988 to 2016, for which we have data from all sites (the plots in Fig. 1 that go back to 1985 are exception, and do not include all sites in the first few years). Plots and analyses (described below) primarily involve species richness or phenological data, specifically dates of first flight (DFF) or dates of last flight (DLF). The later two variables (DFF and DLF) involved filtering to avoid biases associated with variation in the intensity or timing of site visits. Specifically, DFF values were only used if they were proceeded by an absence; in other words, there must be a negative observation before a positive observation is taken as a DFF record. Similarly, DLF values were not used if they were not followed by an absence; so any species observed on the last visit to a site in a particular year did not have a record of DLF for that year. If a species was only observed on a single day in a particular year, then that date was used as a DFF (and only if proceeded by an absence) but not as a DLF, in order to not use the same data point twice.

For a subset of years and sites, absolute counts of individual butterflies (by species) were taken in addition to the presence/absence data; this was done at the 5 lowest elevation sites starting in 1999. These data are used here to investigate the dynamics of the low elevation butterflies during the drought years, specifically the relationship between phenological shifts, changing abundance and dependence on irrigation. For the latter (dependence on irrigation), one of us (AMS) ranked species *a priori* (without knowing the results of analyses) based on natural history observations. Dependency on irrigated areas was categorized as follows: 1), butterfly species that are essentially independent of irrigation; 2), species that use irrigated, non-native hosts in some areas as well as native, non-irrigated hosts in other areas; 3), species that use irrigated, non-native hosts in at least one of multiple generations; and 4), species that are completely dependent on irrigated, non-native hosts.

### Weather variables

Analyses included the following weather variables: maximum and minimum daily temperatures, total precipitation, and a sea surface temperature variable associated with regional conditions^15^. Following previous analyses^13^, maximum and minimum temperatures were averaged and precipitation was totaled from the start of September of the previous year through August of the current year. Previous fall through current summer is a useful climatological time period in a Mediterranean climate and captures precipitation and overwintering conditions that potentially affect butterflies through both direct effects on juvenile and adult stages, and indirect effects through host and nectar plants. Data were generated as monthly values using the PRISM system (Parameter-elevation Relationships on Independent Slopes Model, PRISM Climate Group; http://prism.oregonstate.edu) for latitude and longitude coordinates at the center of each survey route.

As a complement to the site-specific, PRISM-derived weather variables, we used an index of sea-surface temperature associated with the El Niño Southern Oscillation (ENSO). Specifically, we used the ONI (Oceanic Niño Index) values for December, January and February (a single value is reported for those winter months; http://cpc.ncep.noaa.gov) from the winter preceding butterfly observations for a given year. Higher values of this index correspond to regionally warmer and wetter conditions. We also downloaded snowfall data from the Central Sierra Snow Lab located near our Donner Pass site (station number 428, http://wcc.sc.egov.usda.gov/nwcc), but preliminary investigations found that annual snowfall totals were highly correlated with annual precipitation totals. Correlation coefficients between snowfall and precipitation were between 0.80 and 0.88, and the inclusion of snowfall caused variance inflation factors from linear models (described below) to often exceed 10; thus snowfall was not included in final models. In contrast, correlations among other weather variables (maximum temperatures, minimum temperatures, precipitation, and ENSO values) were generally lower: across sites and weather variables, the mean of the absolute value of correlation coefficients was 0.31 (standard deviation = 0.23).

Weather variables that were included in models were *z*-standardized within sites to be in units of standard deviations. This allows variables from sites with different average conditions (e.g., mountain and valley sites) to be readily compared and, more important, it allows for slopes from multiple regression models to be compared among weather variables that are themselves measured on different scales (as is the case with precipitation and temperature).

### Analyses

Statistical investigations involved two phases. First, we used plots of *z*-standardized data to visualize patterns in phenology (DFF and DLF) and flight days; the latter variable, the number of days flying, was expressed as the fraction of days that a species is observed divided by the number of visits to a site per year (this has been referred to as “day positives” in other publications using these data^15^). DFF, DLF, and flight days were z-standardized within species at individual sites and then averaged across species to facilitate comparisons of patterns across sites and years. We also used plots of species richness to explore faunal changes over time at each site. Plots of richness were accompanied by splines (with 5 degrees of freedom) and predicted values from random forest analyses^26^, which both allow for visualization of non-linear relationships. The spline analysis has the advantage of producing smoothed relationships (between richness and years), while the random forest analysis, performed with the randomForest^27^ package in R^28^, has the complementary advantage of being able to incorporate covariates (in this case the number of visits per year) as well as the advantage of making no assumptions about the shape of the relationship (between richness and years, while controlling for sampling effort). Patterns in species richness over time were also explored using diversity indices and Hill numbers that weight rare and common species differently (at different levels of q, which determines the sensitivity of the analysis to rare species)^29–31^, using the vegetarian (v1.2) package^32^ in R^28^. In addition, we used the combination of spline and random forest analyses to investigate changes in abundance (numbers of individuals observed per year) at the low elevation sites where abundance data was available. As with other variables, abundance values were *z*-standardized within species and sites, and *z*-scores were averaged across species.

Following the visualization phase of investigation, we developed simple linear models that were focused on prediction of dates of first flight (DFF) and species richness. Independent variables for both sets of models included average daily minimum temperatures, average daily maximum temperatures, total precipitation, ENSO (ONI sea surface temperatures), sampling effort (number of visits), and year. Models of species richness included date of first flight as an additional variable because we were interested in the possibility that the timing of species emergence affects butterfly populations and consequently observed species richness. These models (for both DFF and richness) were estimated for each site individually and for high and low sites as groups of sites (5 sites in each model). Additional model complexities were explored that included interactions between weather variables, time lags (effects of previous years on current year dynamics) and cumulative effects (sliding windows of averaged precipitation values). Interactions were rare, but we report interactions between weather variables that were significant (at *P* < 0.05). Lagged and cumulative weather variables did not add to the explanatory power of models and the individual lagged and cumulative effects were rarely significant and not discussed further.

As we have found elsewhere for analyses of phenology and richness with these data^11,33^, linear models satisfied assumptions of normality and homogeneity of variance. To address potential collinearity among predictor variables, variance inflation factors were investigated and found generally to be between 0 and 5, and in a few cases between 5 and 10. For instances where inflation factors approached 10, quality control was conducted by including and excluding correlated variables to verify that estimated β coefficients were not affected. Linear models were also used to test the hypothesis that phenological shifts at low elevations have demographic consequences for individual species. For each species at the lowest sites (SM, WS, NS, and RC), we separately regressed dates of first flight against years, and annual abundance against years. Slopes from those regressions were then compared using correlation to ask if species that shifted to an earlier flight (negative slopes for DFF versus years) were also species that become more abundant (positive slopes of abundance versus years). This was done for the years 2008 – 2016 to capture the transition into the millennium drought years, and only included species that were present in at least 6 of those years. As with other analyses, linear models were performed and assumptions investigated using R^28^.

## Acknowledgements

Research was funded by the National Science Foundation (DEB-1638773 to CCN, DEB-1638922 to JAF, and DEB-1638793 to MLF), and MLF was supported by a Trevor James McMinn professorship.

## Author contributions

A.M.S. designed and carried out data collection. J.H.T. and D.P.W. managed data entry and curation. M.L.F., J.A.F. and C.C.N. analyzed the data. All authors contributed to writing.

## Additional information

Supplementary information is available in the online version of the paper.

